# Discovering Differences and Similarities among Species Based on Numeric Features of microRNAs

**DOI:** 10.1101/767046

**Authors:** Rongsheng Zhu, Zhanguo Zhang, Dawei Xin, Yang Li, Qingshan Chen

**Affiliations:** College of Science, Northeast Agricultural University, Harbin 150030, China; College of Agronomy, Northeast Agricultural University, Harbin 150030, China

**Keywords:** microRNA, minimum free energy, numerical feature, different map, species, evolution

## Abstract

Numeric features of microRNA (miRNA) are different from the other RNAs and play a key role in the course of miRNA recognition. Are there significant differences about such numeric features between different species? Are there some species specific about it? In order to answer questions, we implemented the Kolmogorov-Smirnov test for 32 species based on 132 numeric features of miRNAs. Results demonstrate that almost all kinds of miRNA secondary structures matching frequencies show highly similar, and this means that such secondary structure tend to be specific to miRNAs. Length of pre-miRNA, minimal free energy (MFE), and number of stacks show bigger difference between different species, and this means that such features tend to be species-specific. In order to discover differences and similarities among species based on numeric features of miRNAs, we design two tools-species difference map and feature difference map. By species difference map, we find that numeric features of miRNAs can represent class attribute of species and the map basically describe relationship between different species. By feature difference map, we find that there are huge difference about difference of every numeric feature between different species, and this means that strength of such difference is not uniform. Meantime, it means that numeric feature of miRNAs can represent different attributes of corresponding species. Our study present relationship between different species by brand new style-numeric features of miRNAs.

## Introduction

MicroRNAs are endogenous, non-coding, and small-molecule RNAs with a mature length of about 21-23nt. MicroRNAs play an important post transcriptional regulatory role^1^. Their essential functions in cell identity and fate, developmental timing, apoptosis, carcinogenesis, and response to environmental stresses, including disease, has meant that more and more attention has been focused on microRNAs in recent years^2,3,4,5,6,7,8,9,10^. Since the first microRNA was found in *C. elegans* in 1993^11^, 28645 microRNAs had been identified and included in the mirBASE database^12^.

The initial microRNAs were obtained experimentally^13, 14, 15^. Although the experimental results were reliable, it was very difficult to identify those with low expression or those that were only express in a few organs or specific developmental stages^16^. Therefore a new method for identifying microRNAs was required. Statistical recognition technology was introduced to overcome these difficulties^17^, and a large number of microRNAs have been identified and confirmed using this method in recent years^18, 19, 20, 21, 22, 23^.

For the statistical recognition of microRNA, the numerical features of microRNA are the core elements. Minimum free energy (MFE)^24^, based on the minimum free energy algorithm of RNA secondary structure prediction, has become a particularly powerful piece of evidence for discriminating microRNAs from other RNAs^25^. The adjusted minimum free energy (AMFE)^26^ and the minimum free energy index (MFEI)^27^ were derived from MFE and have features that are more specific to microRNAs, and are widely used^28,29^. Research had shown that the secondary structures of microRNAs are highly conserved^30, 31^; therefore, it was used to describe the characteristics of secondary structure paired frequency in the computational identification process^32^. The base content and the entropy of a microRNA are used to describe the microRNA’s base composition characteristics. However, a number of studies^28, 29, 33^ showed that there were obvious differences in the origin, function, base composition, and structure among different species. For example, animal and the plant microRNAs differ greatly in their method of production, mode of action on target genes, enzymes involved, and in the splicing process. Interestingly, the microRNAs showed a high degree of consistency within animals or plants, respectively. We hypothesized that the differences and similarities of microRNAs among species or species class could be depicted by the numerical features of the microRNAs. Thus, the present study aimed to determine whether the numerical features of microRNAs are species- or class-specific and which features have obviously sequence-specific differences or similarities.

Experimental techniques and computational identification methods are being continuously improved, and more and more microRNA genes have been discovered. By end of October, 2010, a total of 15 172 microRNA genes were integrated into the miBASE database^12^, covering nearly a hundred species, and many species were represented by more than a hundred microRNAs. These provided an adequate sample size for the statistical analysis of microRNAs among species. In addition, to fully exploit the numerical features to obtain more comprehensive information on microRNAs, 132 numerical features in eight classes were selected to construct an Eigen vector, which provided a diverse background for our research.

For the analysis method, we selected the Kolmogorov-Smirnov test^34^, a method with higher sensitivity than ANOVA. It can be used to test the distribution difference between two independent samples. To minimize the impact of sampling on the analysis results, a core strategy for our research was designed on the basis of the bootstrap method^35, 36^ combined with the Kolmogorov-Smirnov test.

In the present, the numerical features of microRNAs from 32 species were evaluated by the bootstrap and Kolmogorov-Smirnov tests. The results were used to define a new measure, feature recognition number, to describe the differences among species and to construct a species difference map. Furthermore, another new measure, species similarity degree, was defined to describe the genetic relationship among species and to construct a species relationship map. The results demonstrated that the relationships gained from microRNA prediction were identical with the biological relationships. Our study forms a new method for studying the evolution of microRNAs and species relationships.

## Materials and Methods

### Data Preparation

#### Choice of species

Thirty-two species that had over 100 microRNAs in the mirBASE (Version 21)^12^ were selected, which covered animals, plants, and viruses. All microRNAs from viruses were considered as the same class. In addition, two classes were chosen to verify the research results, one was primates, which included *Homo sapiens, Macaca mulatta, Pan troglodytes*, and *Pongo pygmaeus*, and the other was dicotyledonous plants, which included *Arabidopsis thaliana, Glycine max, Medicago truncatula, Populus trichocarpa*, and *Vitis vinifera*. The basic information on the microRNAs of the candidate species is shown in Table 1.

#### Feature selection

One hundred and thirty-two numerical features in eight classes were selected in total; the serial numbers and names of the features are described in supplementary table 1.

#### Frequency characteristics of single nucleotides

The four features included in the frequency characteristics of single nucleotide were A, C, G, and U. This class is marked as A.

#### Frequency characteristics of dual nucleotides

All the two base combinations of the four bases, A, C, G, and U, were included, such as AA, AC, GU. In total, there were 16 dual-nucleotide features. This class is marked as B.

#### Frequency characteristics of triple nucleotide

All the three base combinations of the four bases, A, C, G, and U, were included, such as AAA, AAG, CGU. In total, there were 64 triple-nucleotide features. This class is marked as C.

#### The frequency characteristics of the secondary structure matching state

Based on RNA secondary structure predicted by Mfold (Zuker^24^), the matching state of each nucleotide was described by the method presented by Xue et al^32^. For example, “A+.” indicates that the nucleotide at the site is “A”, with a non–matching left site, and a non-matching right site in the secondary structure. Examples are shown in Fig. 1. In total, there were 32 frequency features of the secondary structure matching state. This class is marked as D.

**Figure 1.**
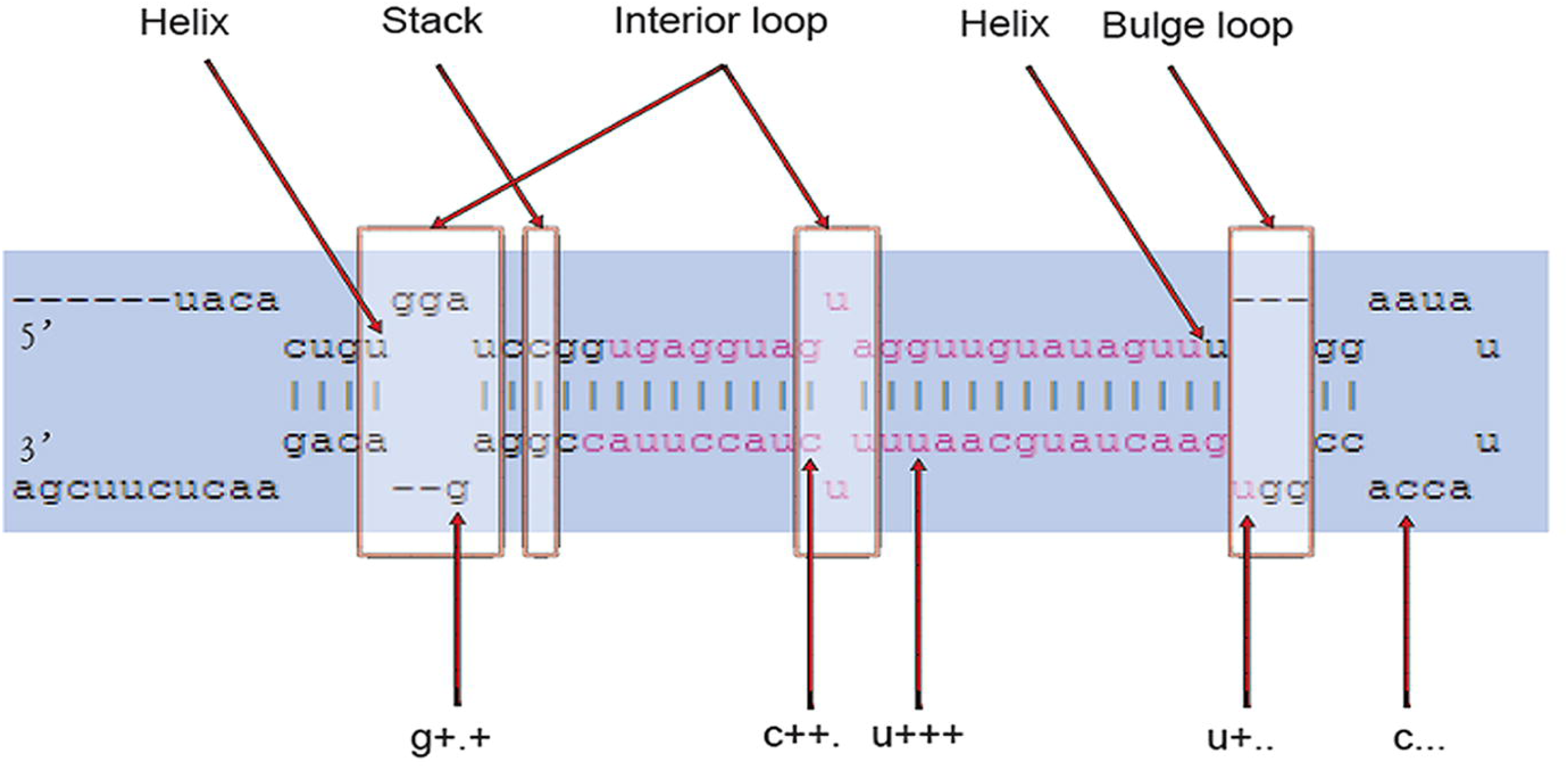
Secondary structure of cel-let-7 as predicted by Mfold. Labels above structure indicate four features of miRNA cel-cet-7: helix, stack, interior, and bulge. Symbols beneath the structure are examples of the secondary structure paired frequency features. For example, ‘g+.+’ indicates that the location base ‘g’ that is not paired, the base to the left of it is paired and the base to the right of it is also paired.

#### Length and the direct count characteristics of the secondary structure

These features included the length of microRNA gene, the number of bulge loops, the number of helices, the number of interior loops, and the number of stacks. Except for the length of the gene, the features were taken from Mfold predictions of the secondary structure. Detailed examples are shown in Figure 1. This class is marked as E.

#### Numerical characteristics related to minimum free energy

This section included the minimum free energy (MFE)^24^, the adjusted minimum free energy (AMFE)^25^, and the minimum free energy index MFEI^26^. This class is marked as F.

#### Base content and base content ratio

The section included G+C content, (G+C)/(A+U) ratio, A/C ratio, and G/U ratio. This class is marked as G.

#### Entropy features

The information entropy^37,38^ was calculated using the formula:

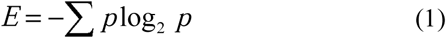

Formula (1) generated four kinds of entropy information related to the frequency of single nucleotides, dual nucleotides, triple nucleotides, and the matching state frequency of the secondary structure (termed Sec_str_entr n supplementary table 1). This class is marked as H.

### Basic analysis method

#### Kolmogorov-Smirnov test

The differences between two samples are usually analyzed by ANOVA (analysis of variance) or by the rank test. However, ANOVA is only used to test the mean of samples, and the rank test requires a similar-distribution assumption between two samples. Thus, a more sensitive and limitless distribution method, the Kolmogorov-Smirnov test^34^, was chosen to test the distribution difference between two independent samples.

Kolmogorov-Smirnov’s test formula is:

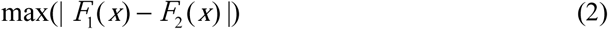

In the formula (2), *F*_1_ (*x*) is the empirical distribution function of the first sample and *F*_2_ (*x*) is the empirical distribution function of the second sample. When the statistic was less than a certain threshold, the null hypothesis was accepted that these two samples had the same distribution; otherwise, the null hypothesis was denied and a distribution difference did exist between these two samples.

#### Bootstrap method

The bootstrap method is a statistical resampling method, and was introduced by Efron in 1979^35^. It is an effective non-parametric test, which is executed by repeated random sampling from original data. In this study, the data the data was subjected to Bootstrapping first followed by the Kolmogorov-Smirnov’s test to obtain more reliable inferences^36^.

#### Research strategy

Basic research strategy is shown in Fig. 2 with an example. *Ciona intestinalis* and *Xenopus tropicalis* were selected as target species and the base content of “A” was selected as a candidate feature. The Kolmogorov-Smirnov test was carried out for each of the 1000 pairs of samples from 1000 bootstrap samplings with a threshold of p = 0.01. The distribution of p-values is shown in Fig. 3(a). When the number was less than the threshold, there was no significant difference between the species, otherwise, the difference was accepted. In our research, the threshold was set at 0.99. In this example, all the p-values from the 1000 tests exceeded 0.01 and the level of the overall test achieved 100% (Fig. 3(a)); therefore, there was significant difference between *Ciona intestinalis* and *Xenopus tropicalis* on A base content and the result was denoted with 1. In Fig. 3(b), most of the p-values from the Kolmogorov-Smirnov test were greater than 0.01 between *Ciona intestinalis* and *Caenorhabditis elegans* on the A base content by 1000 bootstrap samplings and 1000 tests. The overall level of testing (proportion of significant numbers in 1000 tests) was far less than 0.99. Therefore, there was no significant difference between the two species on A base content. The result was denoted with 0.

**Figure 2.**
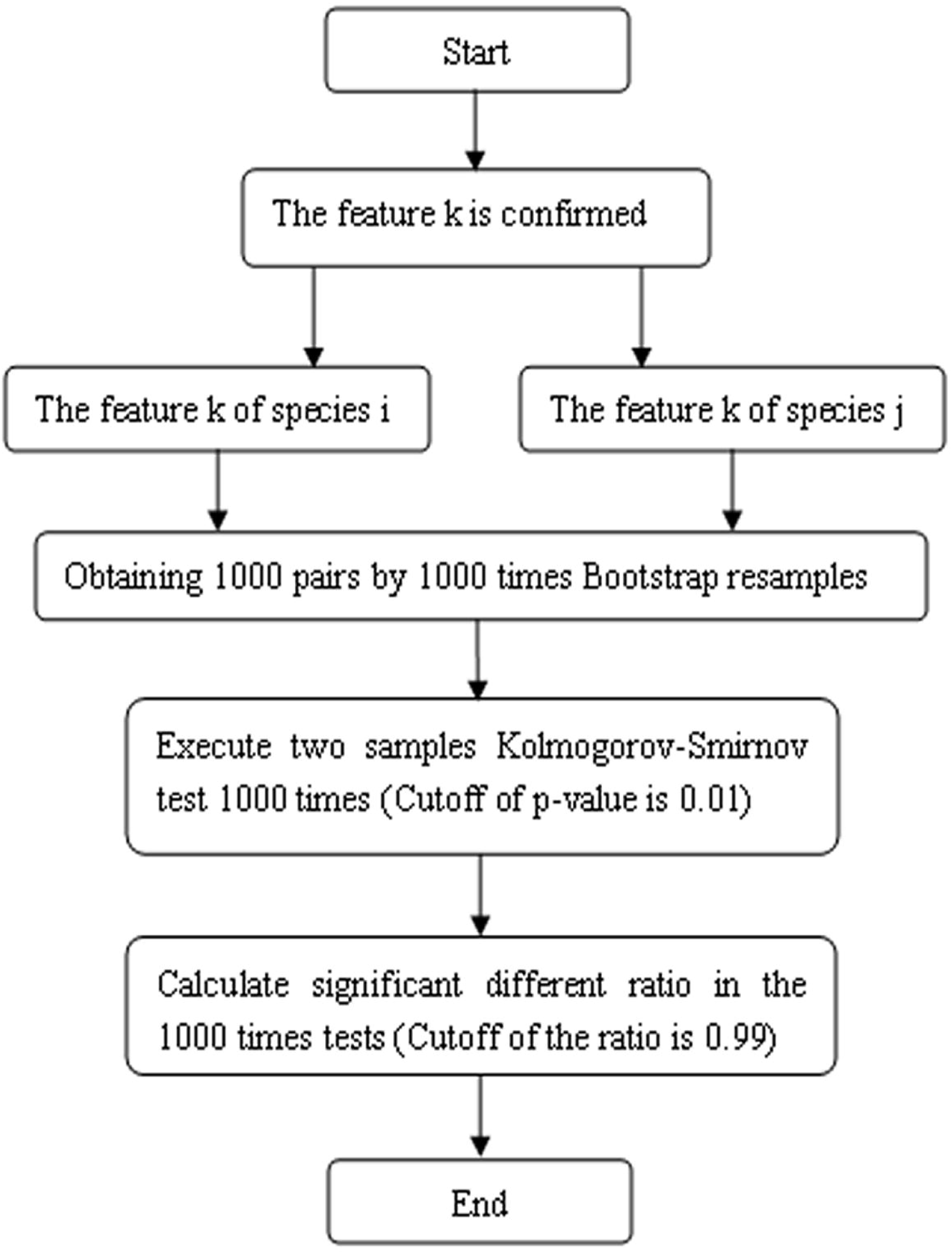
Flowchart of the research strategy. Initially, 132 features were selected, in the flowchart, feature k for species i and j is used as an example. In the second step, 1000 bootstrap resamples were performed and 1000 Kolmogorov-Smirnov tests were executed. In the last step, we calculated significant differences for feature k between the two species, and then made a decision on the species pairs using the significant difference ratio.

**Figure 3.**
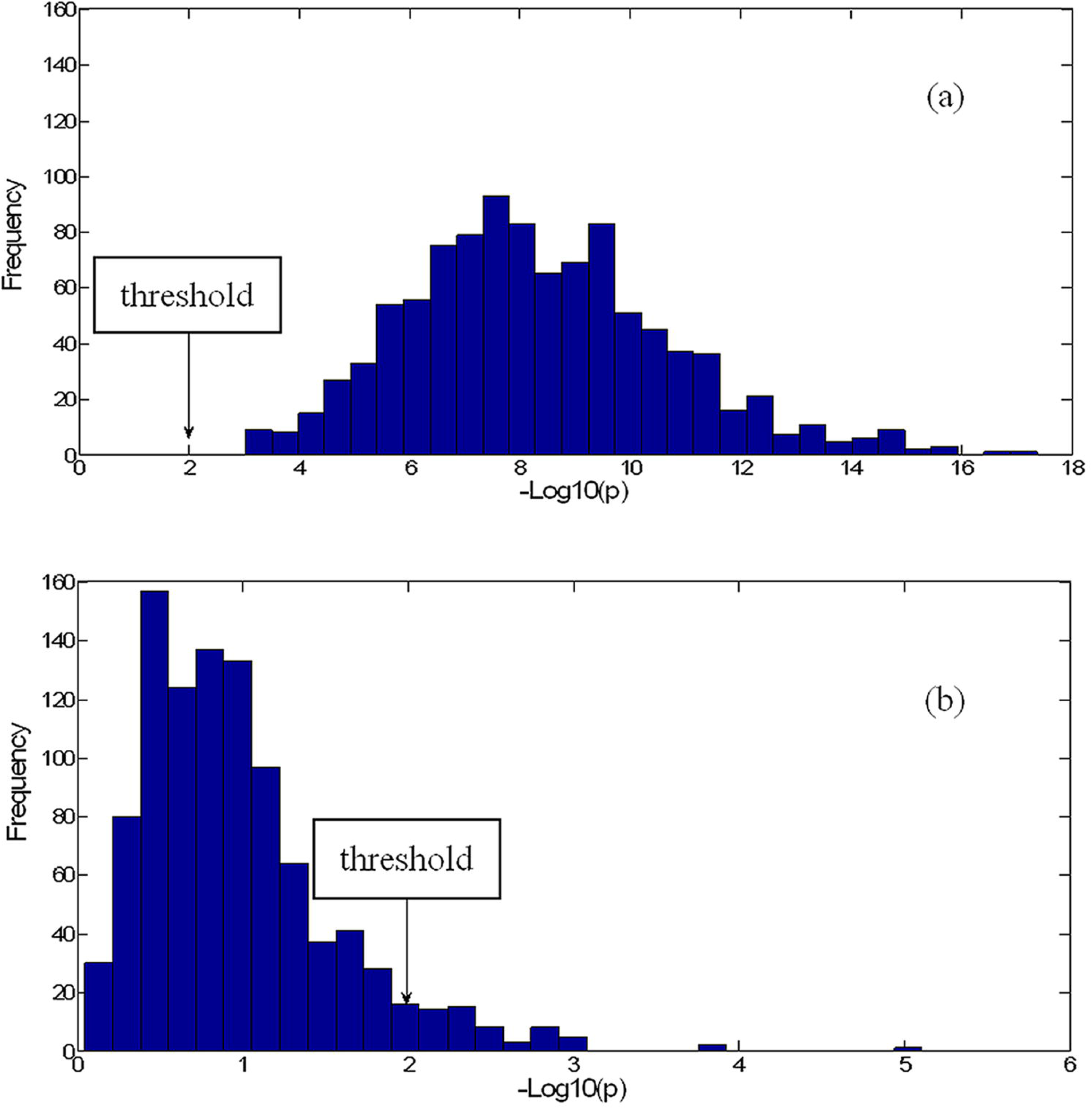
Examples being corresponding to acceptance or refusal of test. (a) Histogram of 1000 K-S tests of the A content of *Ciona intestinalis* and *Xenopus tropicalis* based on 1000 bootstrap values. (b) Histogram of 1000 K-S tests of the A content of *Ciona intestinalis* and *Caenorhabditis elegans* based on 1000 bootstrap values. The cutoff p-value was 0.01.

#### Feature identification number of species pairs (FIN)

Feature identification number of species pairs was the number in candidate features that could be used to distinguish a certain species pair, i.e. there was a significant difference between the species pairs for a certain feature according the basic research strategy. The formula of feature identification ratio (FIR) of species pairs was defined as follows:

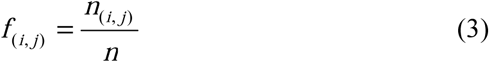

In the formula (3), the *n* denoted the total of candidate features, and the *n*_(*i.j*)_ denoted the feature identification number between species *i* and species *j*.

In this study, FIN and FIR are used to measure the degree of genetic relationship between species.

#### Species pairs identification number of feature (SPIN)

Species pair identification number of feature refers to the number of species pairs that have been distinguished by a certain feature. Subsequently, the species pair identification ratio of feature (SPIR) was defined as follows:

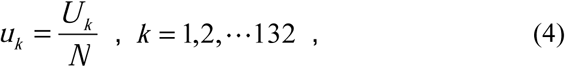

In formula (4), *k* denotes sequence number of the feature, *U*_*k*_ denotes the SPIN value for feature *k*, and *N* denotes total number of species pairs. SPIR could describe the ability of a feature to identify a species in a background of species pairs and was used to evaluate candidate features.

## Results

### Effect of numerical features on species identification

According to the research strategy (Fig. 2), the process should be carried out 496 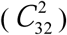 times for each feature. Thus for all 132 features, the process was computed 132 × 496 times. The effect of the numerical features in species identification was obtained from 496 species pairs. The details are shown in supplementary Table 1, and the basic analysis is shown in Fig. 4.

**Figure 4.**
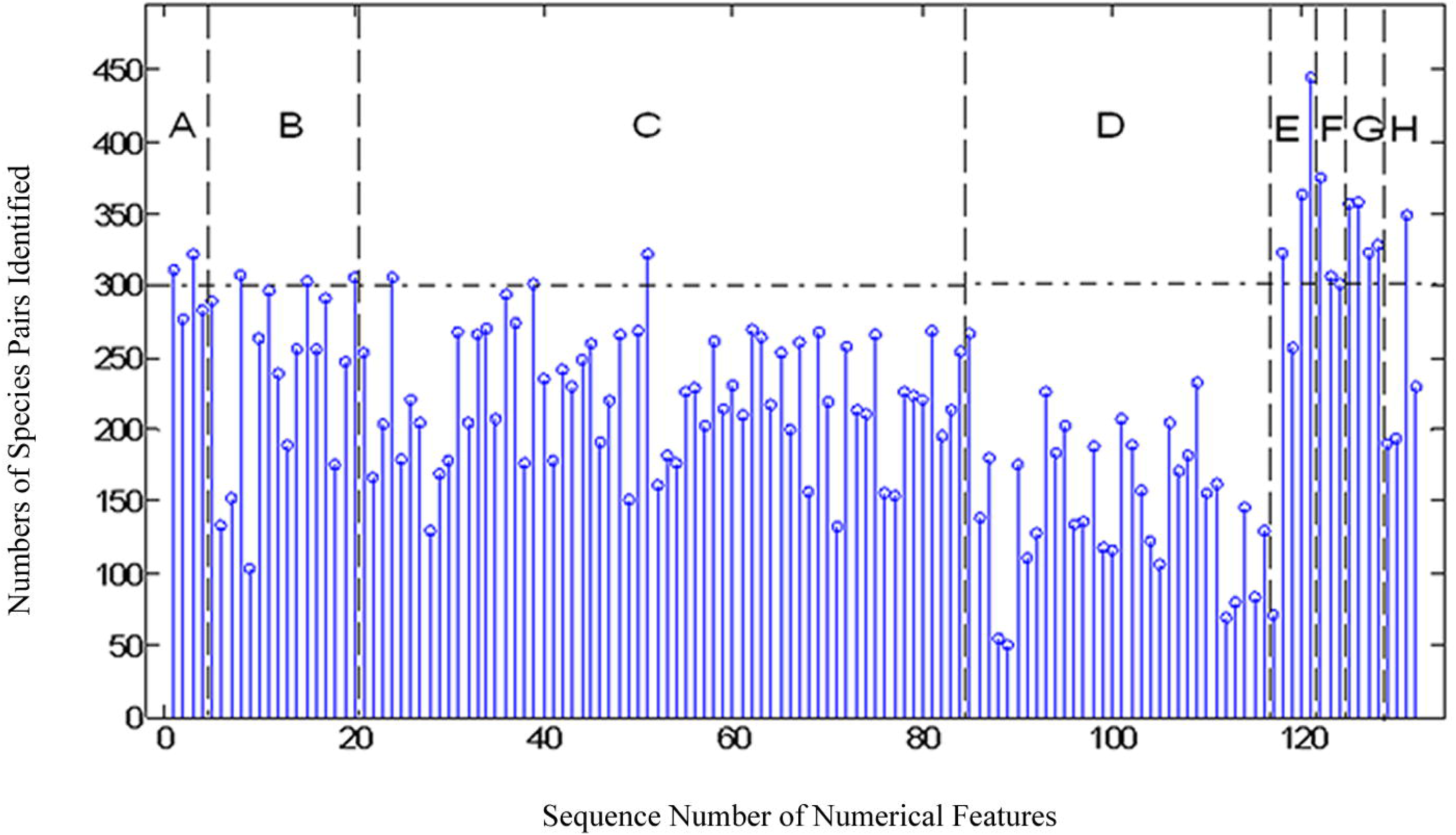
Numbers distribution of species pairs identified in 496 pairs of species based on 132 numerical features. The figure is divided into eight regions (A-H). The regions are described in S_Table 1.

The eight classes of features (A-H) are from left to right in Fig. 4. For the four single nucleotides (class A), the identification ratio among the features was consistent, and the average identification ratio was 60.13%. For the sixteen dual nucleotide features (class B), the identification ratio among the features showed significant differences, and the average identification ratio was 47.99%. For the 64 triple nucleotide features (class C), the identification ratios among the features were moderately consistent, and the average identification ratio was 45.01%. For the thirty-two secondary structure pairs state features (class D), the feature identification ratio was lower than for other classes, and the average identification ratio was 30.31%. This result demonstrated that there was no significant difference among the secondary structures of the microRNAs from all species. For the length and direct counting number of secondary structures (class E), the feature identification ratio of length reached 89.72%, which was the highest of all the features. The feature identification ratios of number of stacks, number of helices, number of interior loops, and number of bulge loops were 73.39%, 65.12%, 51.81%, and 14.31%, respectively. The results showed that there was generally a difference in the length of microRNA genes among species, but there was difference in the number of the number of bulge loops. For the features related to the minimum free energy (class F), the feature identification ratios of MFE, AMFE, and MFEI were 75.6%, 61.9%, and 60.69%, respectively, which showed a higher identification ratio, and a bigger change of features among species. For base content and base content ratios (class G), a higher average identification ratio of 68.85% was observed. For the four entropy-related features (class H), the feature identification of the entropy of triple-nucleotide was 70.36%, the highest in region H, and was less than 50% for the other features

Overall, the top six for recognition rate among the selected 132 features were the sequence length (Length, 89.72% recognition rate), the minimum free energy (MFE, 75.60%), the number of stacks (Stack, 73.39%), (G + C) / (A + U) (72.18%), G + C (71.96%), and the entropy of triple nucleotide content (70.36%). There were significant differences among species identified with these features, and these features were specific for species. However, the bottom six features for recognition rate were A.++ (10.08%), A +.. (11.08%), U +.. (13.91%), Bulge loop number (14.31%), U .++ (16.13%), and U ++. (16.94). There was little difference among species identified with these features; however, they were more specific among microRNAs.

### Identification of species pairs based on SPIN

Four hundred and ninety-six species pairs were identified with the 132 numerical features, and the results are shown in Supplemental Table 2 and Fig. 5.

**Figure 5.**
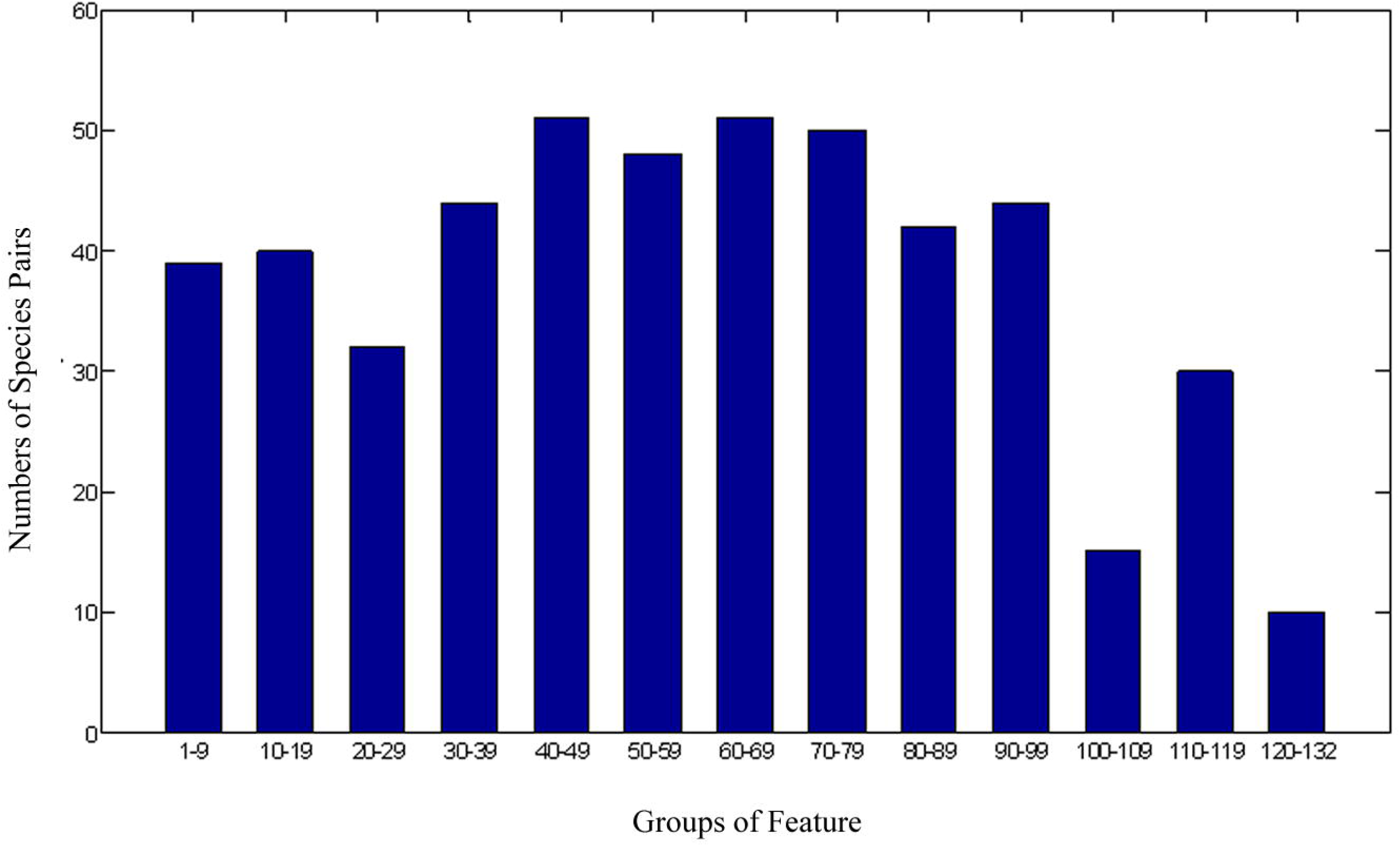
Distribution of numbers of species pairs on the basis of identification feature numbers.

In Fig. 5, 39 species pairs were distinguished by no more than 10 features, and 10 species pairs could be distinguished by more than 120 features.

A genetic relationship map among species was constructed with the criterion that SPIN was no more than 10 between any two species (Fig.6). Number 19 (*Drosophila pseudoobscura*) and number 20 (*Caenorhabditis elegans*) were in same class of *Ecdysozoa*. Number 27 (*Populus trichocarpa*), number 28 (*Vitis vinifera*), number 30 (Sorghum bicolor), and number 31 (*Zea mays*) were in another class of *Magnoliophyta*. Furthermore, a genetic relationship should exist among species from number 2 to number 17, which are all vertebrata. The SPIN among some species pairs, including (7,8), (7,9), (7,10), (8,9), (8,10), and (9,10), were 2, 0, 0, 3, 4, and 0, respectively, which showed a high similarity degree among the microRNAs of primates.

**Figure 6.**
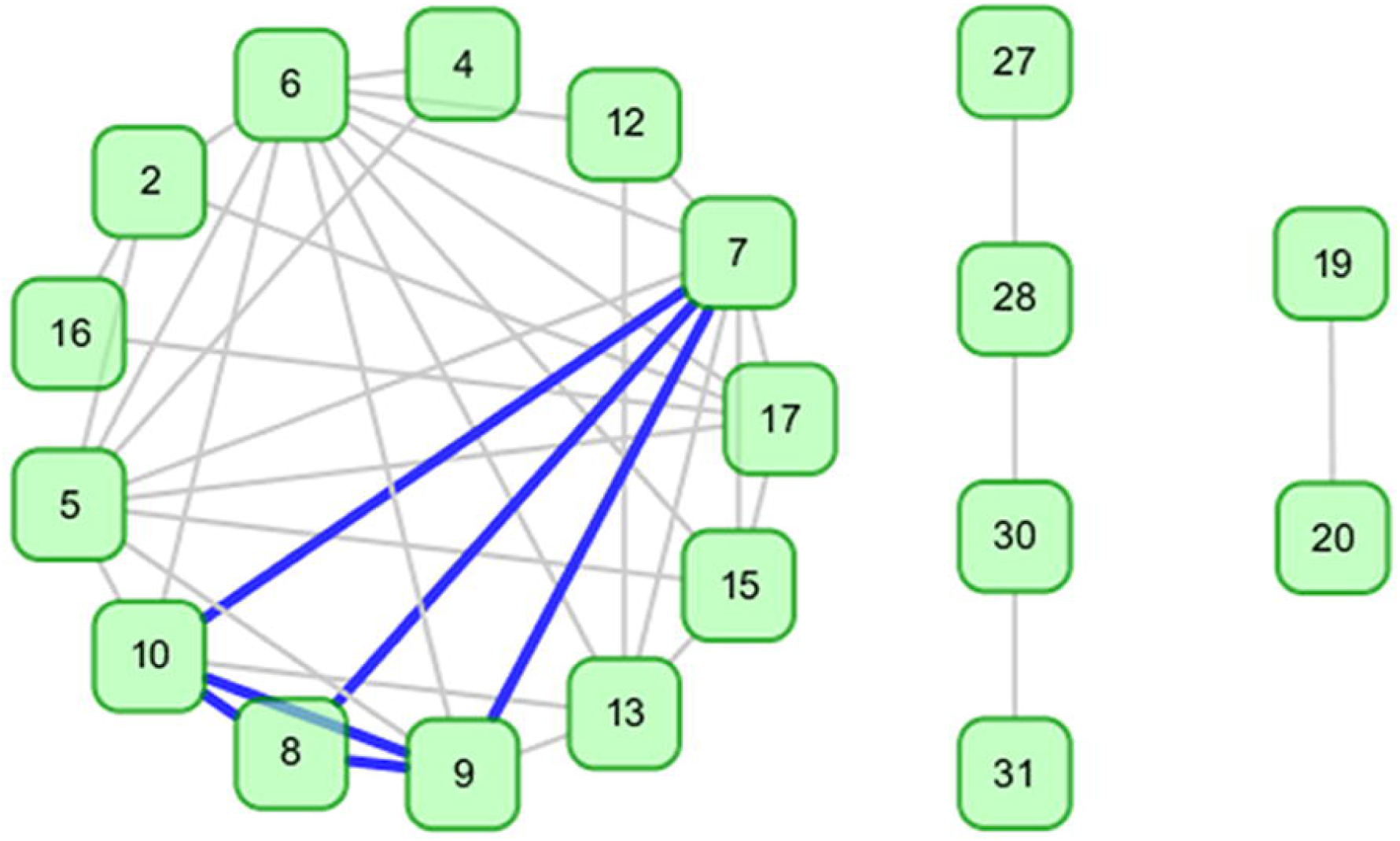
Graph of species relationship. Graph of species relation based on species pairs difference feature numbers that are less than 9. The numbers in the figure represent the candidate species: 2-Xenopus tropicalis, 4-Canis familiaris, 5-Equus caballus, 6-Monodelphis domestica, 7-Macaca mulatta, 8-Homo sapiens, 9-Pan troglodytes, 10-Pongo pygmaeus, 12-Mus musculus, 13-Rattus norvegicus, 15-Sus scrofa, 16-Danio rerio, 19-Drosophila pseudoobscura, 20-Caenorhabditis elegans, 27-Populus trichocarpa, 28-Vitis vinifera, 30-Sorghum bicolor, 31-Zea mays.

There were ten species pairs, (1,23), (4,24), (4,26), (8,24), (8,26), (12,24), (12,26), (14,24), (14,26) and (26,32), whose SPIN values exceeded 120. The first nine pairs are all species pairs between animals and plants. The last one, (26,32) is a species pair between a plant and a virus. The results showed that there was a large difference between plant and animal microRNAs. However, no significant difference was found between animal and virus microRNAs. Ultimately, there should be fewer different features between species with closer genetic relationships.

In Fig. 5, except for the species pairs in either end of the spectrum, which were obviously very similar or very different, respectively, most of the species pairs could not be used as a complete standard to explain the relationship between the two species. To describe the state of species pairs distinguished by features, a difference map among species was constructed based on 132 numerical features from 32 species.

### Species difference map

A 32×32 matrix was constructed among 32 species from the species classification order. The definitive element of the matrix was the SPIN between two corresponding species. The matrix was named the species difference matrix (SDM), on the base of numerical features of microRNAs. Furthermore, based on the SDM, a hot map was drawn as a difference map of species pairs or species difference map (SDMP) (Fig. 7).

**Figure 7.**
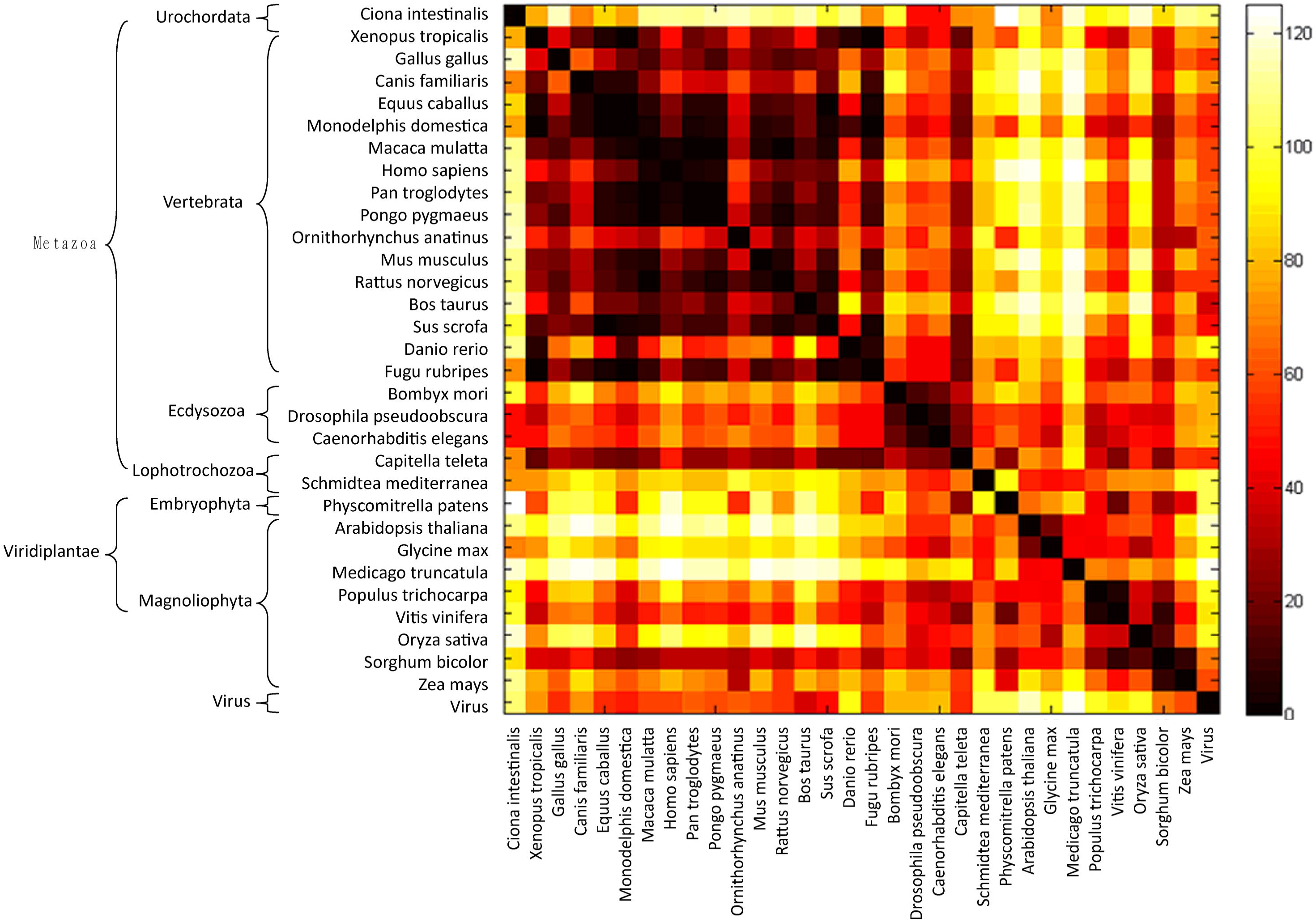
A hot map demonstrating the state of 32 species by number of feature identified among 132 features.

From Fig. 7, the dark square regions in the upper-left and lower-right corners indicated that there were few differences in microRNA features among animals or plants. Moreover, the upper-right and lower-left corners of the map have brighter colors, which indicate that there was a clear difference in terms of microRNA features between animals and plants. The greater the genetic distance among species, the brighter the color of the elements in the matrix, which provided strong evidence that SPIN values are able to describe the genetic relationships of species pairs.

There were very few differences between the *Ciona intestinalis* from Urochordata and *Bombyx mori, Drosophila pseudoobscura*, and *Caenorhabditis elegans* from Lophotrochozoa; however, there were more differences between *Ciona intestinalis* and the other species, which indicated that there was a closer genetic relationship between Urochordata and Lophotrochozoa than that with either of them and the other species. Similarly, there were clearly dark squares among the three animal species in Lophotrochozoa, indicating a high similarity among species in the same class, but there were fewer SPINs between the species of the Lophotrochozoa and *Xenopus tropicalis, Danio rerio*, or *Fugu rubripes* from Vertebrata, and *Capitella teleta* from Lophotrochozoa.

In the sixteen species from vertebrata, there was apparently a difference between *Danio rerio* and the other species, and the SPIN values among the other species, except *Danio rerio*, were very small, which showed that microRNA genes are highly conserved among vertebrata species and among animal species as a whole. Furthermore, a smaller SPIN value was obtained between *Danio rerio* and *Fugu rubripes*, than with other species, which showed that there was high conservation within those species.

For plants, the matrix was darker among the species in Magnoliophyta, but brighter between Magnoliophyta and the other classes. The extent of the color was apparently inferior to that of vertebrata. The results showed that there were bigger differences among the microRNA genes of species in Magnoliophyta, which supported the hypothesis that there is a large diversity in microRNA genes of plants. The matrix also shows the viral microRNA genes are more similar to those of animals than plants.

In summary, the SDMP constructed by SPIN could directly display the degree of difference between species pairs. The SDMP could display the basic differences and similarities between species, classes, or within classes. Thus SDMP could be used as a valid tool to display the genetic relationship of species.

### Relationship of species based on similarity degree of identification (SID)

SPIN could effectively reveal the genetic relationships among species. However, SPIN or SPIR showed the reverse changes with extent of genetic relationship of species. Therefore, the Similarity degree of identification (SID) was calculated as a consistent measure of the genetic relationship of species. SID was defined as follows:

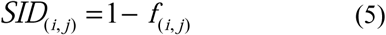

In formula (5), *i* and *j* denote the sequence number of the species. The *f*(*i, j*) denotes the SPIR values of species *i* and species *j*. From the definition of SID, the bigger the value of SID, the closer the genetic relationship between the species. Based on this measure, a relationship map of all candidate species was constructed (Fig. 8).

**Figure 8.**
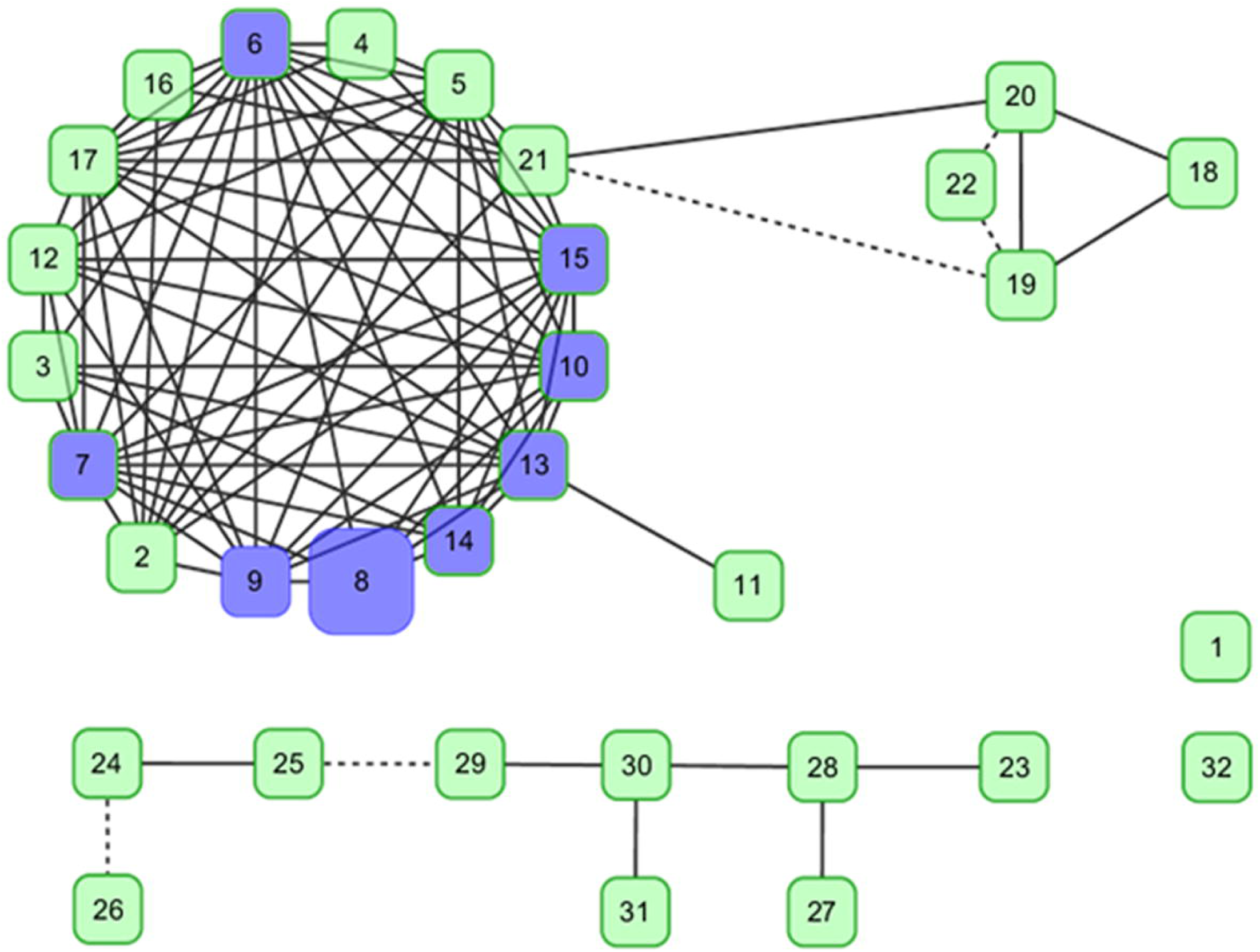
Graph of species relationship. 32 species’ relation graph on the base of similarity degree of identification (SID) with a cutoff of 0.8. The solid line indicates a SID of more than 0.84 and the dotted line indicates a SID between 0.8 and 0.84. The number eight represents Homo sapiens and those species that link with it directly are colored blue, some of which belong to the primates. The numbers in the figure refer to the candidate species: 1-Ciona intestinalis, 2-Xenopus tropicalis, 3-Gallus gallus, 4-Canis familiaris, 5-Equus caballus, 6-Monodelphis domestica, 7-Macaca mulatta, 8-Homo sapiens, 9-Pan troglodytes, 10-Pongo pygmaeus, 11-Ornithorhynchus anatinus, 12-Mus musculus, 13-Rattus norvegicus, 14-Bos taurus, 15-Sus scrofa, 16-Danio rerio, 17-Fugu rubripes, 18-Bombyx mori, 19-Drosophila pseudoobscura, 20-Caenorhabditis elegans, 21-Capitella teleta, 22-Schmidtea mediterranea, 23-Physcomitrella patens, 24-Arabidopsis thaliana, 25-Glycine max, 26-Medicago truncatula, 27-Populus trichocarpa, 28-Vitis vinifera, 29-Oryza sativa, 30-Sorghum bicolor, 31-Zea mays, 32-Virus.

The threshold of SID was set as 0.8. The candidate species were divided into four classes, animal, plant, virus, and Urochordata, respectively. In figure 8, *Bombyx mori, Drosophila pseudoobscura, Caenorhabditis elegans, Capitella teleta*, and *Schmidtea mediterranea*, who have similar shape or habit, were classified into the same group by SID analysis. All primates were classified into the same group as *Homo sapiens. Glycine max, Arabidopsis thaliana*, and *Medicago truncatula*, belonging to Magnoliopsida, were classified into the same sub-class; and *Oryza sativa, Sorghum bicolor*, and *Zea mays*, belonging to Liliopsida, were classified into another sub-class. These results showed that SID could reflect the genetic relationships among species.

### Features difference map (FDM)

SPIN or SID were both able to comprehensively evaluate genetic relationships for a pair of species based on their background features information. However, they involved counting numbers of different features between species pairs, and could not display the extent of the difference for every feature between species pairs. To assess the detailed differences and similarities between species for a certain feature, a feature difference map (FDM) was designed, based the numerical features of microRNAs.

The FDM is a gray map, and indicates the extent of the genetic relationship among 32 candidate species. The FDM was built using statistics of species difference calculated by Kolmogorov-Smirnov test (see Fig. 2) with a certain features of microRNAs.

Eight features (representing classes A-H) were selected from the 132 candidate numerical features of microRNAs to illustrate the FDM. The eight features were the C base frequency, AA base pairs frequency, AAA base frequency, A …frequency, number of helices, MFE, G+C content, and entropy of ternary bases. The eight FDMs were built by the average of 1000 Kolmogorov-Smirnov tests with 1000 bootstrap resamples from original sample of species pairs. The results are shown in Fig. 9.

**Figure 9.**
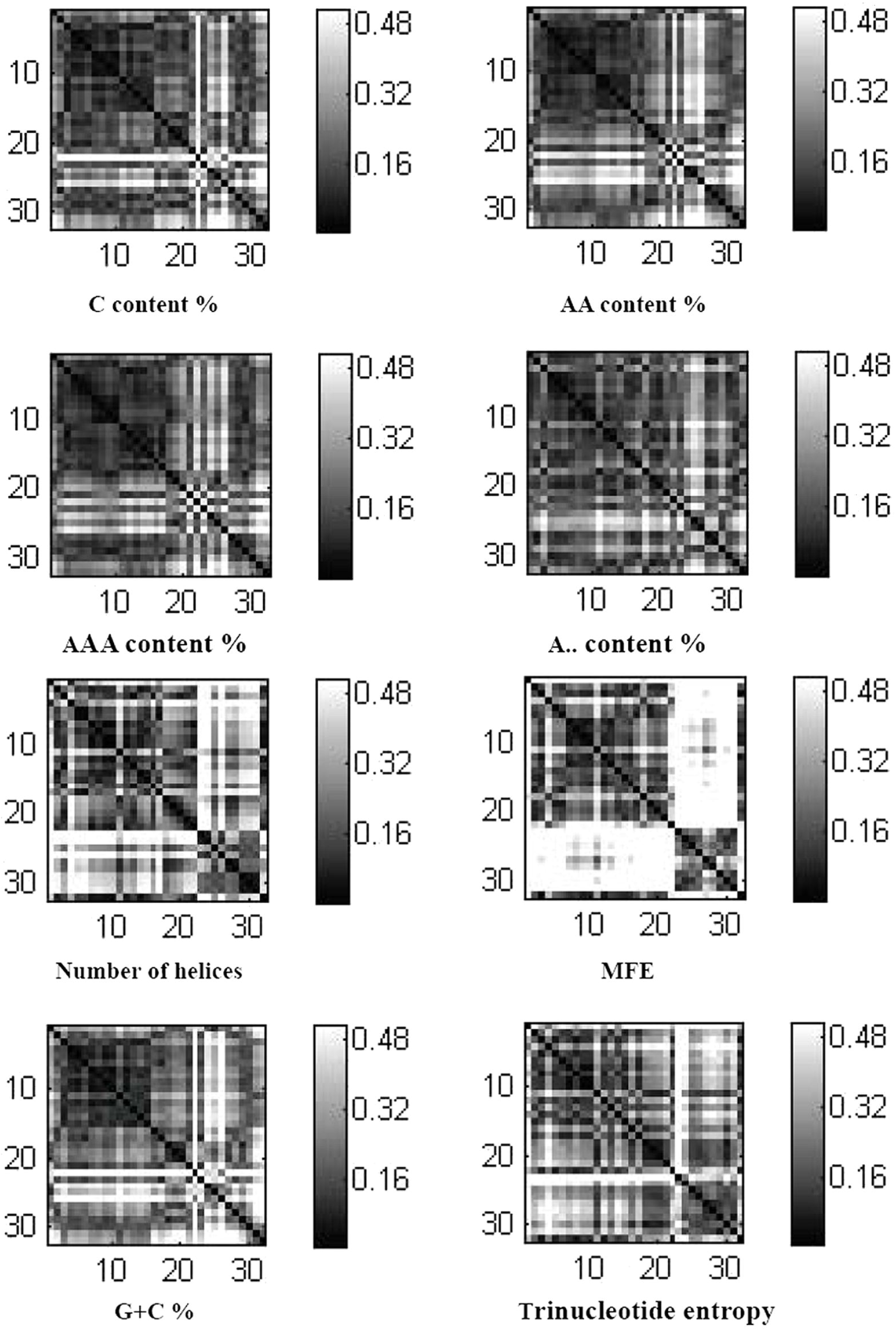
Examples of feature difference map. Eight subplots showing the distribution differences for eight features of the 32 species by Kolmogorov-Smirnov test statistics. The eight features represent the eight regions shown in Figure 4 and detailed in S_Table 1.

The eight FDMs illustrate the relationship among species based on the similarities and differences for each feature. The FDM could be considered as supplementary to SDMP, and it was beneficial for discovering the essential differences between species.

In Fig. 9, the classification indicated in the eight FDMs are roughly consistent with the biological classes, indicating that the Kolmogorov-Smirnov test statistic could be used as an measure of the extent of similarities and differences between species or classes. There were some differences among the eight FDMS in Fig. 9. The four FDMs in the top row are less distinct than the other four in the bottom row. The SPIRs of the four in the bottom row were higher than those of the four top row, with the SPIR of the clearest MFE reaching 75.6%. This result demonstrated, to some extent, the ability of SPIR to distinguish species by microRNA numerical features. The differences between the eight FDMs could be explained by the one-sided relationship of species described by a single feature.

To describe the general differences and similarities between species, we propose that the SPIN value should be used. Of course, the SPIN can only describe the relationship between species generally, because extent of differences between the same species pairs varied for the different features. However, it is reasonable to hypothesize that the SPIN could describe the extent of the differences between species under an effective background provided by a large number of numerical features of microRNAs.

## Discussion

Almost all kinds of biology functional molecular have specific nucleotide composition or special molecular structure^39, 40, 41,42,43,44^. Certainly, miRNAs is the same. Plenty of research projects illustrate that miRNAs is a class ancient biology functional small molecular and is considered as important evolution tag^45, 46,47,48,49,50,51,52^. Lots of jobs have been finished based on sequence analysis of miRNAs^50, 53, 54, 55^. From new perspective, we analyze specificity of miRNAs with relationship to species based on their numerical signal. By our analysis, we have inferred three hypothesis as following.

### The miRNAs numerical signal owned its selection as facing different species

By FIR analysis, lots of evidences are found. For example, FIR of A.++ is 0.1008, but FIR of C… is 0.4556. It demonstrate that there are more different in C… than A.++ among species and corresponding analysis result is shown in S_Fig 1. From the figure, C… content percentage is bigger variation than A.++ content percentage among species. It indicate that secondary structure of miRNAs is more sensitive to C… content percentage not A.++ content percentage. The same evidence is number of bugle loop and length of pre-miRNA and analysis results are shown in S_Fig 2. The finding is that length of pre-miRNA is more sensitive than number of bugle loop of miRNA molecular. Interestingly, if the number of bugle loop of miRNAs is too much, the secondary structure of miRNAs will become unstable and miRNAs will not be able to perform its duties. So it analayze, we find that number of bugle loop is from 0 to 7 and 90 percent of miRNAs is from 0 to 3. Analysis results are shown in S_Fig 3. Less number of bugle loop is corresponding to former researches^56, 57, 58^.

### Species prefer to some numerical features of miRNAs

By SPIR analysis, we find that the SPIR of some species pairs is higher than the others and it indicate that there are more significant different numerical features in those species pairs. For example, the SPIR of Xenopus tropicalis and Monodelphis domestica is 0 and the SPIR of Xenopus tropicalis and Arabidopsis thaliana is 0.7121. It demonstrate that there are no difference between Xenopus tropicalis and Monodelphis domestica and they have close relationship. On the contrary, there are lots of significant different numerical features between Xenopus tropicalis and Arabidopsis thaliana. Certainly, from biology classification, we explain easily the reasons about the results. Because Xenopus tropicalis and Monodelphis domestica both belong to vertebrata, but Arabidopsis thaliana belong to plant. So those differences derive of differences of the species themselves. We apply t-test to both species pairs based on all the numerical features of miRNAs and results are shown in S_Fig 4(Upper figure). The findings demonstrate that some features, which refer to two test consistent non-significant features, show have no significant difference between species and others have significant differences. The non-significant features maybe are miRNAs specific features, but significant features maybe are species specific features. The same test have been implemented in *Canis familiaris*-*Equus caballus* and *Canis familiaris*-*Medicago truncatula* and results are shown in S_Fig 4(Under figure). The same conclusion have been obtained. Our results are similar to former researches^56, 59^.

### Numerical features of miRNAs can dissect relationship and difference among species

In order to describe relationship and difference between species, SID has been designed as measuring species relationship method. Analyzing results are shown in Fig 8. The findings demonstrate that results basically correspond to real species classifications. Furtherly, it indicate that an effective new methods, which can efficiently differ relationship of species, maybe can be created based on miRNAs numerical feature.

## Conclusions

Based on the strategy of the Kolmogorov-Smirnov test combined with the bootstrap method, 132 numerical features of microRNAs from 32 species were analyzed. The results indicated that some features had higher ability to distinguish species, and that other features tended to be conserved between species. In addition, the differences and similarities in numerical features of species; microRNAs reflected the differences and similarities of certain attributes or characteristic of the species to different extents, indirectly indicating the differences and similarities in the shape, habitat, or classification. Furthermore, the genetic relationship among species was analyzed with SPIN, which showed that the closer the genetic relationship between species or within a class, the fewer the differences in features. Subsequently, SDMP were constructed based on the SPIN values. The basic differences between pairs of species and whole differences and similarities of species could be described from the SDMP. Finally, FDMs were designed based on the Kolmogorov-Smirnov test to illustrate the differences and similarities between species or a particular feature. The FDM of the feature was a beneficial supplement to the SDMP.

In this study, we provided a new strategy for using microRNA numerical features to provide insights in the relationship and difference of microRNAs to depict the genetic relationships of species from the perspective of microRNA numerical features.

## Additional Information

### Competing Financial Interests Statement

The authors declare no competing financial interests.

### Author Contribution Statement

Rongzheng Zhu and Qingshan Chen wrote the main manuscript text, Zhanguo Zhang and Yang Li analyzed all data and designed all programs, Dawei Xin and Zhaoming Qi prepared all figures. All authors reviewed the manuscript.

