## Supplementary figures and images for "Discovering Differences and Similarities among Species Based on Numeric Features of microRNAs"

### S_Fig 1

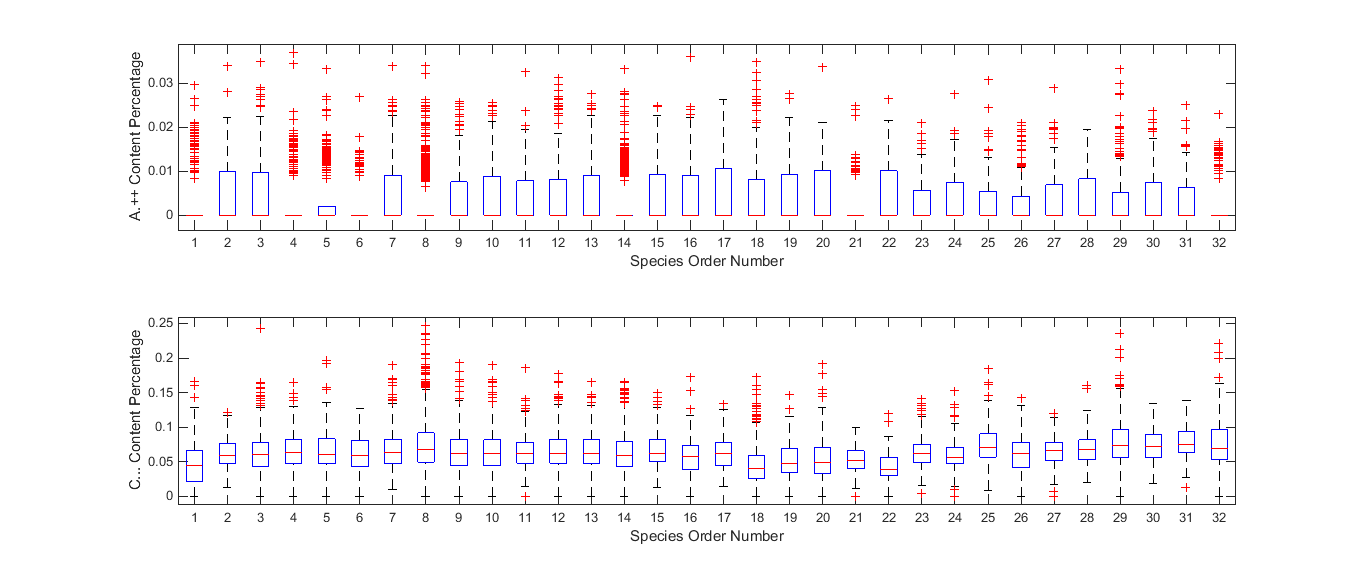

### S_Fig 2

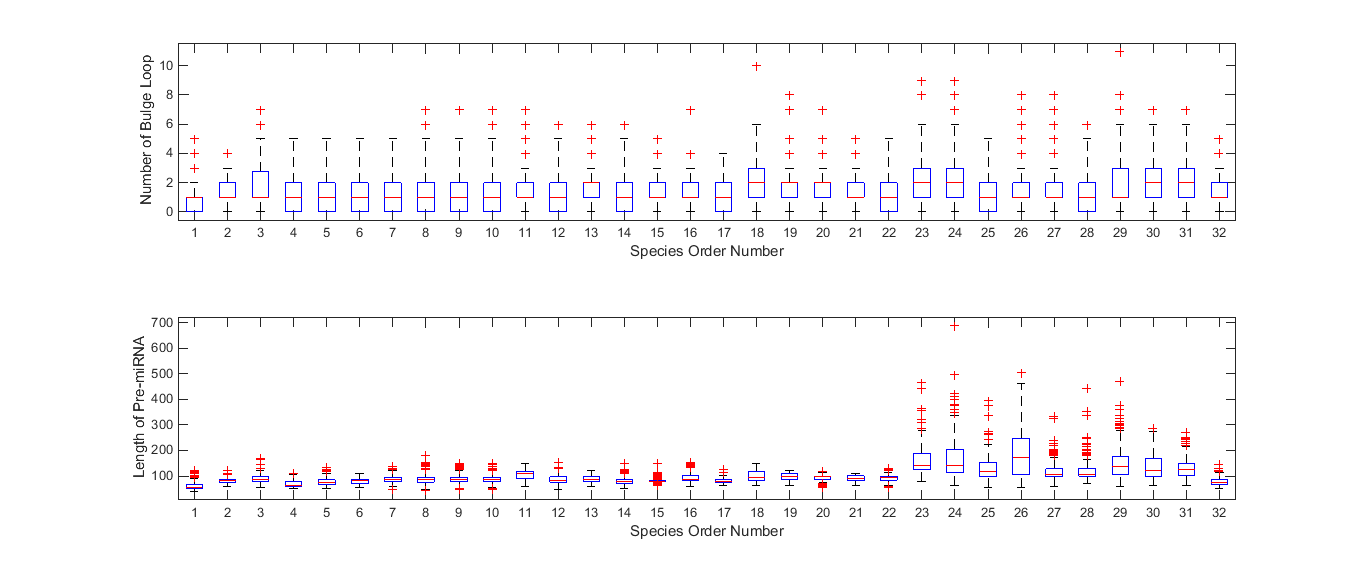

### S_Fig 3

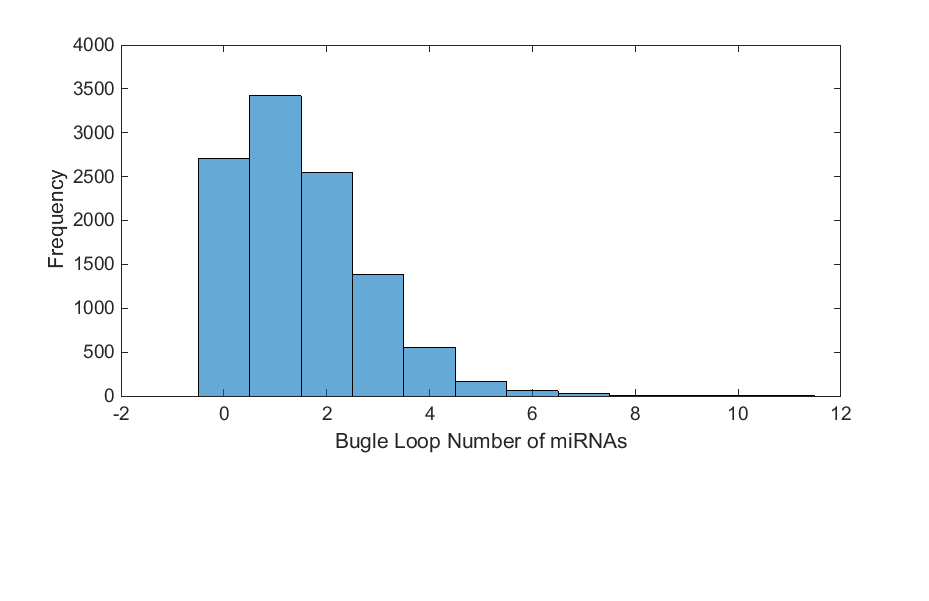

### S_Fig 4

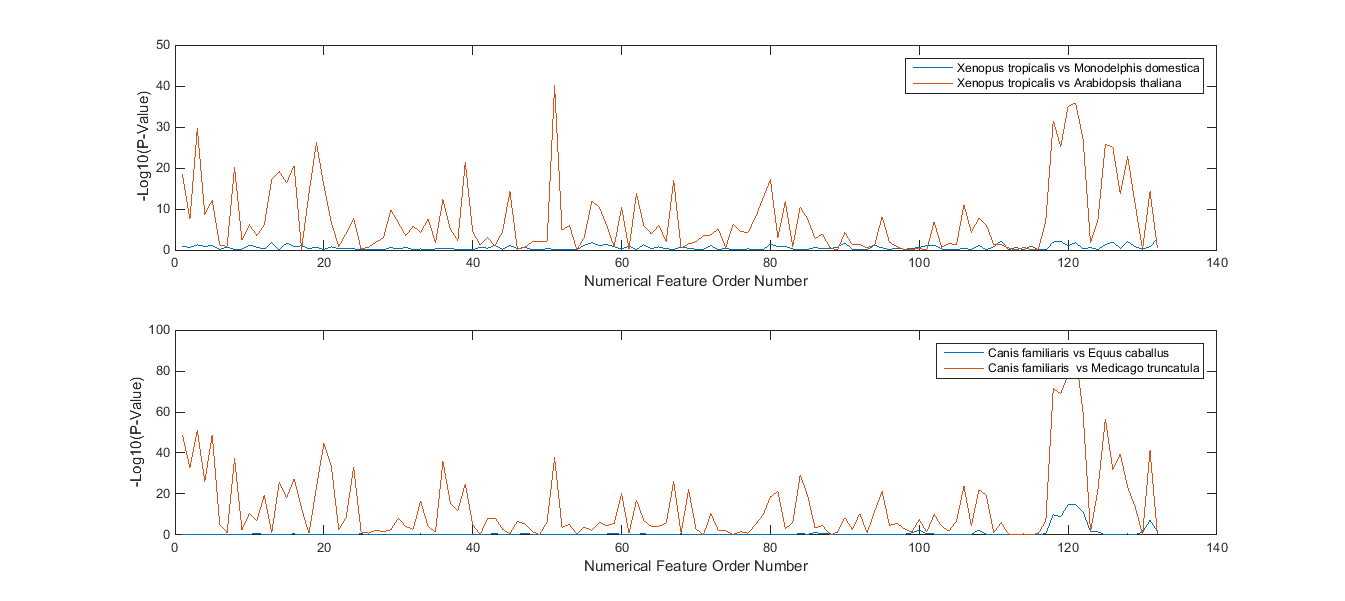
